# Acclimating to degraded environments: The social rationale for swift action on restoration

**DOI:** 10.1101/2022.09.27.509807

**Authors:** Vadim A. Karatayev, Robyn S. Wilson, D.G. Webster, Mark Axelrod, Chris T. Bauch, Madhur Anand

**Author notes:** Conceived study: VAK. Designed model: VAK, RSW, DGW, MA, CTB, MA. Analyzed model: VAK. Wrote paper: VAK, DGW, RSW, MA, CTB.

## Abstract

As environmental degradation progresses, economies and societies adapt to the loss of ecosystem services and public attention to degradation subsides. In systems experiencing such societal acclimation to degradation, net incentives for stakeholder mitigation peak during early degradation phases and subside over time. Using harmful algae blooms in western Lake Erie as a case study, we illustrate how declines in public attention and societal reliance on lake recreation (i.e., finding recreation alternatives) reduce the incentives for stakeholders to reduce pollution runoff (i.e., mitigation efforts throughout the watershed). We then analyze how acclimation can affect a broad array of conservation challenges by developing a general socio-ecological model of societal response to degradation. We find that delays in initiating stakeholder-driven mitigation efforts can exponentially prolong restoration projects. Furthermore, when alleviating intense degradation relies upon voluntary commitments by many individuals, windows of opportunity for mitigation can be very limited because feedback loops of societal adaptation doom late restoration efforts to failure and lock human-environment systems into degraded states. These windows of opportunity can be particularly narrow when a) stakeholder mitigation requires supportive public opinion or b) even modestly valuable alternative services are available in degraded ecosystems. In such cases, maintaining undegraded human-environment regimes may hinge on quickly initiating stakeholder mitigation movements and allocating limited government conservation funds soon after degradation begins instead of spreading mitigation efforts out over decades. Such initiatives, regardless of whether acclimation is slow or rapid in a given system, also greatly accelerate the pace of environmental restoration.

**Significance Statement:** As societies acclimate to degraded environments, mitigation efforts that hinge on action by many stakeholders can erode. Developing a socio-ecological model of acclimation, we reveal how social and environmental processes intertwine to create alternative stable socio-ecological regimes, with either: 1) undegraded ecosystem states sustained by widespread mitigation adoption, or 2) degraded states where societies neither maintain nor continue relying on traditional, local ecosystem services. This dynamic places a premium on prompt mitigation efforts, which may face narrow opportunity windows to get started and avert degraded regimes in systems that rely on stakeholder-driven mitigation. Moreover, in any system requiring stakeholder action, societal acclimation will increase the importance of early action because decaying mitigation incentives exponentially lengthen restoration efforts.

---

**G**iven rapid growth of human population and technology, maintaining undegraded ecosystem states and services often requires mitigating unsustainable anthropogenic impacts. Classical examples of successful conservation involve decisive actions by central actors, for instance changes in resource use regulations to rebuild depleted fish stocks (1, 2) or large government infrastructure projects to reduce pollution (e.g., sewage treatment plants). Insufficient political will to enact regulations and distribute funds can prevent such government responses, however, and many conservation efforts require actions or regulation compliance by many private actors (e.g., reducing fertilizer use on farms to address water quality challenges; 3). Rapid action by a large proportion of stakeholders therefore is increasingly a prerequisite for successfully mitigating environmental degradation and restoring natural ecosystems (4).

Effective collective action for environmental governance can be costly and delayed in most societies. Recognizing environmental crises and identifying effective mitigation strategies takes time (4–6), and mitigation requires individuals’ time and financial resources on top of the costs of reducing harvests or decreasing pollution levels (7). Investing in mitigation can also carry opportunity costs as alternative investment opportunities offer larger or more certain short-term payoffs (‘discounting the future’, 8). From a rational-choice perspective, effective collective action fails when these individual costs outweigh the potential benefits of mitigation to the individual. Critically, when mitigation benefits decline over time, environmental governance movements that begin later also face reduced (and potentially negative) net incentives for stakeholders to act.

Mitigation benefits can decline over time as human societies acclimate to prolonged environmental degradation via three characteristic pathways (Table 1). First, users reduce their reliance on specific ecosystem resources or services. For example, beach use, recreational fisheries, and coastal property values gradually decline as harmful algal blooms become frequent, but people often switch to alternative recreation activities (9–12). Second, societies shift to alternative service that become available as environmental degradation progresses. In many ecosystems, overfishing of predators (e.g., cod) has led to an expansion of prey species (e.g., lobsters) that fishers gradually track with a switch in gear and targeted species (‘fishing down the food web’, 1, 2, 13).

**Table 1.**
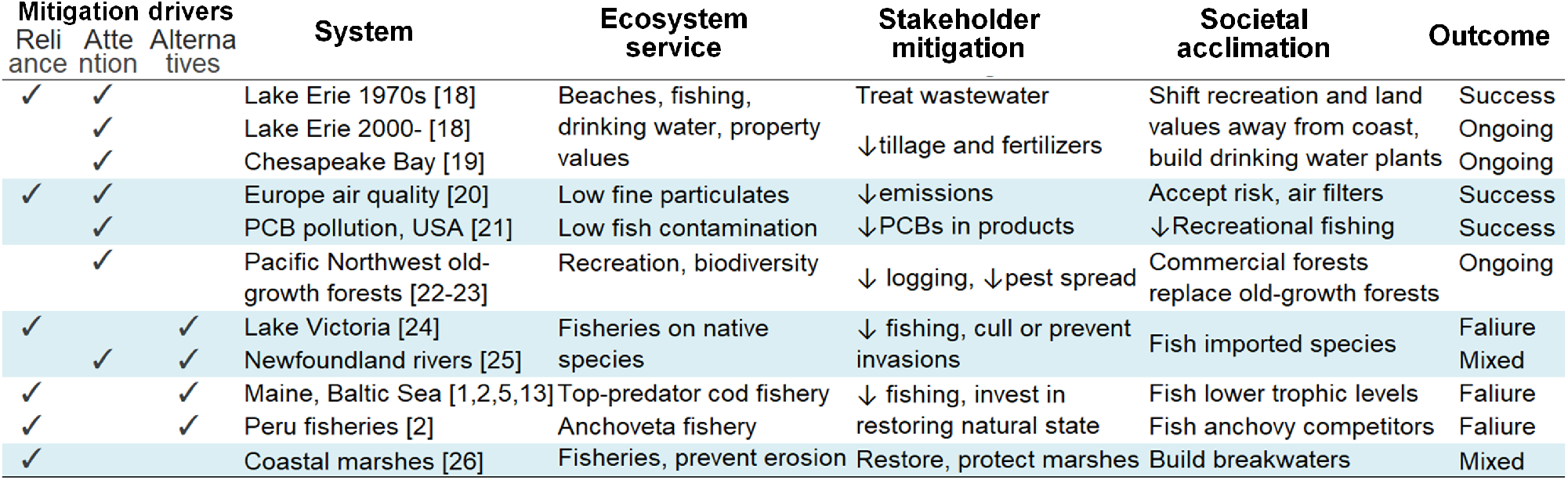
Systems with potential acclimation feedbacks affecting stakeholder-driven mitigation. Mitigation drivers denote the types of acclimation most likely to affect stakeholder mitigation in each system. “Mixed” outcomes denote conservation scenarios where the outcome of conservation is system-dependent.

In addition to shifting away from natural ecosystem services, societies become desensitized to emerging environmental crises. Major events such as wildfires or algal blooms initially gain widespread attention. However, as other issues gradually dominate the discourse, especially when stakeholders realize that addressing the problem is costly, public attention and the impetus for mitigation subsides - completing an “issue-attention cycle” (Fig. 1a; 14–16). These pronounced but limited periods of public attention may be critical when powerful stakeholders anticipate limited economic benefits from restoration. For example, coastal communities who use affected waters for livelihoods or recreation will garner most of the benefits of reducing nutrient pollution, while inland communities still incur mitigation costs (17). Similarly, indigenous communities and others that greatly dependent on a natural resource will benefit most from effective resource management, but roving bandits – resource users who can diversify their resource portfolios – simply switch to exploiting new resource stocks rather than paying high opportunity costs of mitigation driven by reduced harvests (2). Taken together, declining socioeconomic incentives for stakeholder mitigation that delay restoration and prolong resource declines feed back to further reduce mitigation incentives, creating an *acclimation feedback* loop of perpetually diminishing mitigation.

**Fig. 1.**
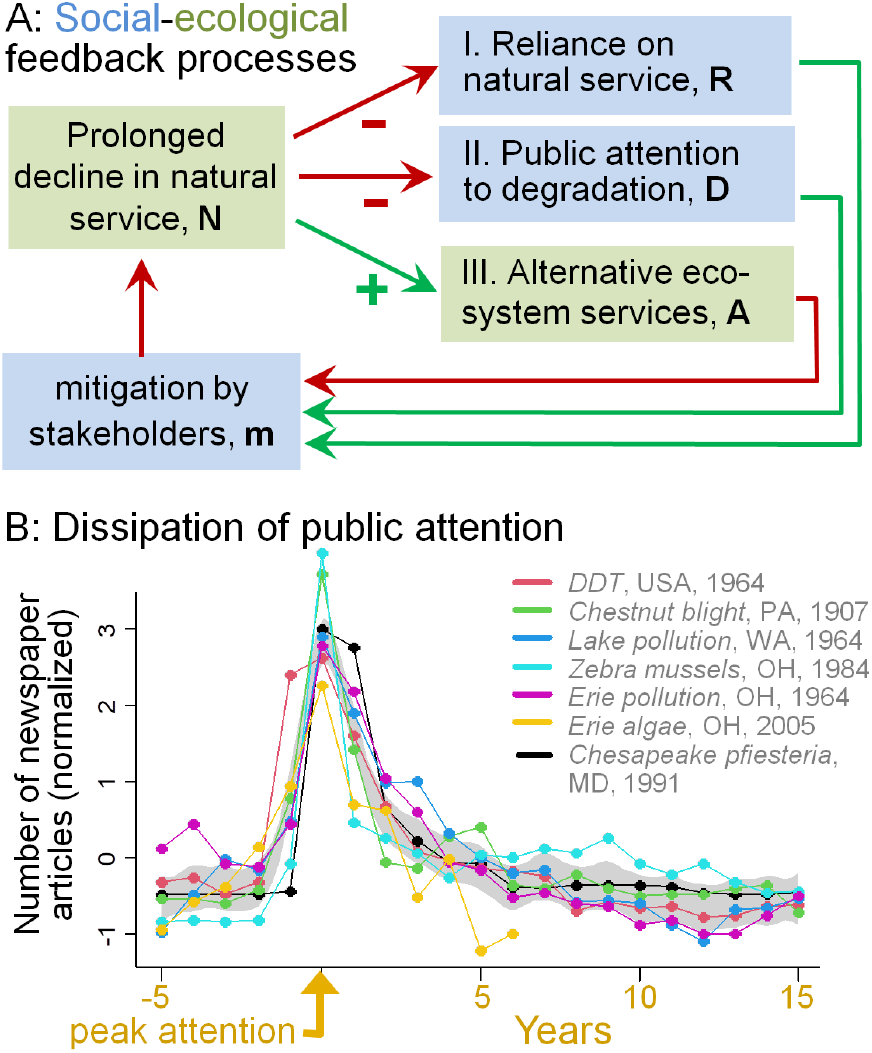
Coupled social-ecological feedbacks affect stakeholder mitigation. (A) Three forms of societal acclimation (I-III) to environmental degradation that can produce feedbacks to impede mitigation; bold letters denote model variables. (B) Public attention, approximated by the normalized number of newspaper articles in a region, can quickly decline from peak levels independently of whether environmental degradation is abated (e.g., lake pollution, DDT) or not (e.g., zebra mussels, chestnut blight). Legend denotes term, area, and start year used in searches on newspapers.com. Shaded area denotes a 95% confidence interval of a spline fitted to all time series.

Given the reality of issue-attention cycles and societal acclimation to environmental change, we pose the question: to what extent can societal acclimation feedbacks reduce the chance and pace of ecosystem restoration? Starting with a general socio-ecological model of ecosystem degradation, we compare how the importance of early action for restoration outcomes differs across three acclimation scenarios: (i) declining reliance on ecosystem services, (ii) declining public attention, and (iii) shifting reliance onto alternative services that emerge in degraded ecosystems (Table 1, Fig. 1a). Next, we explore how the likelihood of successfully restoring undegraded socio-eclogical regimes depends on environmental versus societal processes, and government conservation strategies specifically. We then review how acclimation feedbacks shape conservation in a range of ecosystems (Table 1), and highlight two instances where acclimation has (1) allowed and (2) hindered efforts to reduce pollution in Lake Erie over the past six decades.

## A framework of acclimation to degraded environments

We first develop the scenario where societies acclimate to degradation by reducing reliance *R* on a natural-state ecosystem service *N* (‘natural service’ hereafter), and then extend this model to alternate mechanisms of acclimation. We track the service level as a proportion of its steady state maximum, which changes with ecosystem recovery rate *r_N_* and is being depleted by degradation *μ*. Degradation arises from a fixed amount of stakeholders and can represent, for instance, cod abundance declining with overfishing or lake water quality reduced by organic pollution. Degradation can be reduced to a degree *b* by mitigation, of which a portion *f* depends on the proportion of private stakeholders *m* that elect to mitigate. The remaining proportion of mitigation 1 – *f* is undertaken by the public sector, for example government environmental conservation agencies (‘government mitigation’ hereafter), yielding

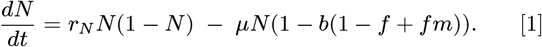

Societal changes in economic activities and recreation habits can lead to changes in natural service reliance *R* that happen at a characteristic rate *ξ*. This rate increases when societies acclimate to the loss of natural services more quickly, for instance when many alternative income, food, or recreation opportunities exist:

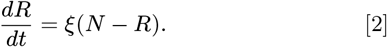

Many conservation efforts rely to a substantial degree *f* on stakeholders in the ecosystem electing to reduce their impacts by following conservation recommendations or regulations. The proportion of stakeholders that mitigate *m* changes at a social learning rate *κ* and depends on the net expected payoff of mitigation Π, yielding

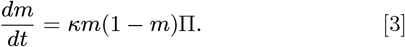

The payoff Π can increase with stakeholder reliance, the value natural-state services *π_N_*, and public influence, and decrease with the presence of alternate services and the cost of mitigation *c_m_*. Throughout, we rescale *π_N_* and *κ* such that *c_m_* = 1.

A second way societies acclimate to degradation is through declines in public attention that favors mitigation. Public attention to degradation often follows a sharp increase as societies become aware of the loss of natural-state services, and subsequently subsides as public focus shifts to other issues (Fig. 1c; 14, 15). We model attention using an epidemic-type dynamic, where individuals gradually progress from (U) unaware of degradation to, (D) aware and actively discussing degradation, and finally to (O) aware of degradation but predominantly discussing other issues. Assuming information spreads with rate *β*, attention shifts to other issues with rate *δ*, and service risk 1 – *N, U* follows *dU/dt* = –*βUD*(1 – *N*) and the proportion of society concerned by degradation follows

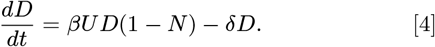

In making mitigation decisions, stakeholders place a weight 1 – *p* on service reliance and a weight *p* on the influence of public attention. Public attention is proportional to the size *s* of the human population exerting attention; for every parameterization we calibrate this parameter such that 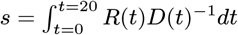. Finally, non-substituability of natural ecosystem services or dedicated conservation movements constantly maintain a proportion 1 – *ω* of mitigation incentives even with prolonged service depletion.

A third way societies can acclimate is by using alternative ecosystem services for which demand gradually expands as a result of natural-state service degradation. This often arises when overexploitation of one species dominant in undegraded ecosystems allows a prey or competitor species to increase in abundance (e.g., ‘fishing down the food web’ 1, 2). We assume that the undegraded state of the ecosystem, represented by the level of *N*, inhibits at a rate *α* alternative ecosystem uses and services. As degradation progresses, alternate ecosystem services *A* have a value *π_A_* and grow at a rate *r_A_*, yielding

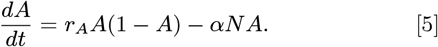

Combining all three sources of acclimation, the net payoff of mitigation for stakeholders is

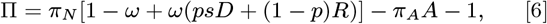

where *π_A_* = 0 is analogous to an absence of altered-state services and *p* = 0 represents no influence of public attention.

### Societal acclimation slows restoration

Does the tendency to acclimate place a large premium on early mitigation efforts in society? We focus on the time (and potentially overall costs) needed to restore an ‘undegraded’ socio-ecological regime (high service *N*, reliance, and mitigation) as a function of time delay *b_t_* between when degradation begins and when stakeholders begin to mitigate (i.e., *κ*(*t*) = 0 for *t* < *t_b_* and *κ*(*t*) = *κ* otherwise). We assume low initial stakeholder mitigation *m*_0_ = 0.05 and that 30% of all mitigation efforts lie in government action and begin immediately at the onset of degradation.

In the base case of reliance-driven stakeholder mitigation (*p* = 0.25, *π_A_* = 0), we find that delays in mitigation exponentially increase the time needed to restore ecosystem states (Fig. 2). This arises because decreasing reliance on the natural-state service reduces incentives for stakeholders to participate in mitigation, especially for lower natural-state service payoffs. Thus a 6-year delay in stakeholder mitigation at *π_N_* = 6, for example, increases restoration time by four decades (Fig. 2a).

**Fig. 2.**
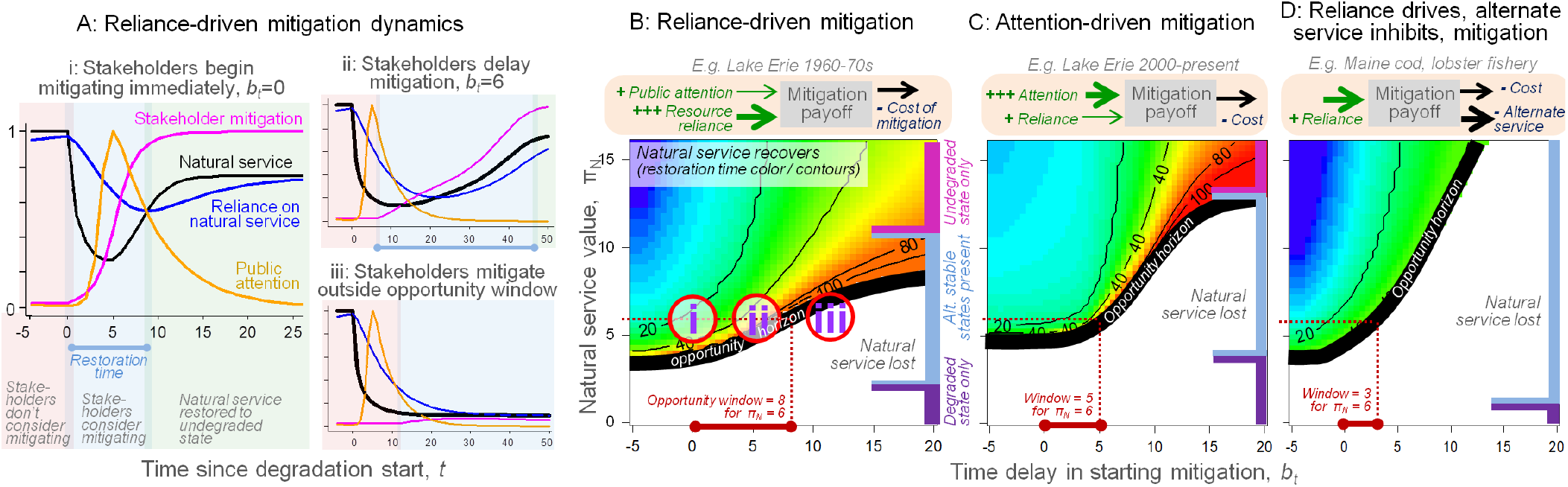
Socio-ecological feedbacks slow the restoration natural ecosystem services and produce limited time windows outside which late mitigation efforts fail. (A) Simulations exemplifying reliance-driven mitigation for three delays (i-iii) between the start of degradation and the time when stakeholders begin to mitigate *b_t_*. Shaded boxes denote regimes before (red) and after (blue) stakeholder mitigation begins, and after the natural-state service is restored (green), with blue bars representing restoration times (restoration fails in iii). Simulations in (A) correspond to letters in (B). (B-D) Mitigation outcomes across values of mitigation delay (*b_t_*, x-axes) and natural-state service value (*π_N_*, y-axes) in scenarios where incentives for mitigation are driven by: reliance on the natural service (B, *p*=0.25, *π_A_*=0), public attention (C, *p*=0.75, *π_A_* =0), or reliance and switching to alternative services (D, *p*=0, *π_A_* >0). Colors and contours measure restoration time and white areas denote conditions under which restoration never occurs. Opportunity window length (maximum delay *b_t_* for which restoration occurs) is the x-axis value that falls on the thick black line for each level of natural service value (exemplified by red bars for *π_N_* =6). In (B-D), blue rectangles denote *π_N_* regimes that lead to limited opportunity windows and alternative stable state dynamics and purple (pink) rectangles denote *π_N_* regimes where only degraded (undegraded) steady-states exist (see Fig. S1); *b_t_* < 0 values represent cases where stakeholder mitigation precedes degradation.

Societal acclimation also slows restoration in model parameterizations of qualitatively different dynamical regimes. Compared to reliance-driven mitigation, the length of time required for successful restoration increases when stakeholder mitigation incentives come primarily from public opinion (75%) rather than reliance (25%; *p* = 0.75, *π_A_* = 0, Fig. 2c). This occurs despite our parameterization of *s* that makes *total* mitigation incentives equal between the reliance- and opinion-driven scenarios. Instead, the lower potential for public opinion to drive mitigation arises because opinion *concentrates* mitigation incentives into a brief period of widespread public attention, after which discourse shifts to other issues (Fig. 1c). Finally, restoration slows further when degradation gives rise to an altered-state service *A*(*p* = 0, *π_A_* > 0, Fig. 2d), for instance when overfishing predators causes an increase in valuable prey species.

### Societal acclimation can limit opportunity windows for restoration

We find that societal acclimation can also change restoration outcomes qualitatively. After prolonged service depletion and delays in mitigation, many individuals no longer rely on the natural-state service, leaving insufficient incentives for stakeholders to commit to mitigation. This feedback loop stabilizes a degraded socio-ecological state where societies do not expect to benefit from, and do not mitigate, a depleted service. As a result, societies can face a limited window of opportunity (*b_t_* values along the black border in Fig. 2b-d), where stakeholder-driven mitigation efforts that begin after this window passes never restore the ecosystem.

When limited opportunity windows exist, their length can be short in many conservation scenarios, such as when natural-state services are less valuable (Fig. 2b) or when mitigation incentives are primarily driven by public attention (Fig. 2c). Very short opportunity windows also arise in the presence of an altered-state service *A* (Fig. 2d), even when the natural service is five times more valuable than the alternate service (*π_R_* > 12.5 *vs. π_A_* = 2.5). This potentially leaves mitigation efforts that preempt degradation (*b_t_* < 0) as the only feasible path to restoration.

Limited windows of opportunity for restoration also signal the presence of underlying alternative stable states, an insight we confirm in Fig. S1. Under this dynamic, undegraded ecosystem regimes are more resilient if they can recover from a larger disturbance. Systems with longer opportunity windows - where undegraded states recover after a longer period of un-mitigated degradation and a greater cumulative disturbance - therefore also have greater ecological resilience (27). We show that this positive window length - resilience relation generalizes to both environmental and social dimensions of the system in Appendix A. Thus, longer opportunity windows correspond to a greater likelihood of avoiding degraded socio-ecological regimes more generally as disturbance events impact both the environment (e.g., natural disasters) and society (e.g., economic recessions reducing stakeholder mitigation).

### Drivers of socio-ecological resilience

Which conservation management interventions might expand opportunity windows and resilience? One approach is to increase the proportion of mitigation conducted by government agencies (1 – *f*) rather than private stakeholders (*f*). However, even when governments conduct 40-60% of mitigation, limited resilience and opportunity windows can arise and delays in stakeholder mitigation *bt* disproportionately protract restoration (Fig. 3a).

**Fig. 3.**
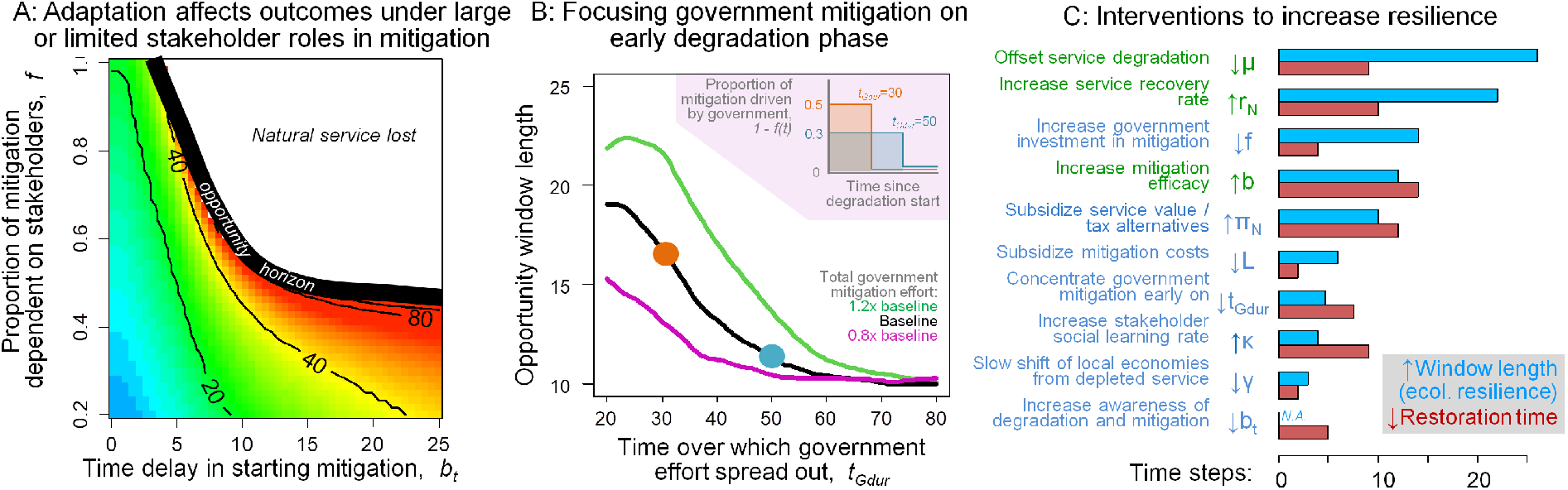
Factors driving the length of opportunity windows and socio-ecological resilience. (A) Acclimation feedbacks can prevent or greatly slow restoration even when a fraction of total mitigation effort depends on stakeholder action. Colors and contours denote restoration time across the range of mitigation delays *b_t_* (x-axis) and the proportion of mitigation that depends on private stakeholders *f* (y-axis); white areas denote conditions under which restoration never occurs. Following Fig. 2b-d, opportunity window length is the x-axis value that falls on the thick black line for each level of *f*. (B) Opportunity windows expand when a fixed amount of total government mitigation is concentrated on a narrower period of time *t_Gdur_* (left to right along x-axis), with lines representing 80% (purple), 100% (black), and 120% (green) of non-stakeholder effort in our base case. Inset in (B) demonstrates how f is varied over time to maintain constant total government effort (1 – *f*)*t_Gdur_* for two levels of *t_Gdur_*. (C) Local sensitivity analysis of the effect of environmental (green) or social (blue) processes on reliance-driven mitigation scenarios. Blue bars represent the decrease in time needed to restore the natural service; red bars represent the increase in opportunity window length (i.e., the maximum time delay *b_t_* for which restoration succeeds). Increases (decreases) represent a change in parameters from 80% to 120% (120% to 80%) of baseline values. Analyses (A-C) correspond to reliance-driven stakeholder mitigation (Fig. 2a-b).

Alternatively, government agencies may have the option to concentrate their efforts between the onset of degradation (*t* = 0) and a time when government mitigation ends *tGdur*, in order to buy time for stakeholder mitigation to expand (i.e., *f*(*t*) = *f* for *t* < *t_Gdur_* and *f*(*t*) = 1 for *t* > *t_Gdur_*). We find that windows of opportunity grow when the same amount of government effort TotEff_*Govt*_ = (1 – *f*)*t_Gdur_* is concentrated on a shorter period of time, and this benefit of early government mitigation increases with TotEff_*Govt*_ (Fig. 3b). Finally, comparing all possible management interventions, we find that societal features impact both resilience and restoration time to a similar order of magnitude as environmental features (Fig. 3c). This suggests that informed policy can compensate for environmental features that erode resilience in some systems.

## Case study: Lake Erie pollution

The processes underpinning limited opportunity windows (Fig. 1a) exemplify the history of pollution control in western Lake Erie (Fig. 4). This socio-ecological system has faced two qualitatively different restoration efforts: (1) 1960-70s reliance-driven mitigation of coastal pollution with a moderate role of elective stakeholder action *f* (*sensu* Fig. 2b, 3a) and (2) contemporary attention-driven mitigation of watershed pollution that hinges on voluntary participation by farmers with low reliance on lake water quality (*sensu* Fig. 2c).

**Fig. 4.**
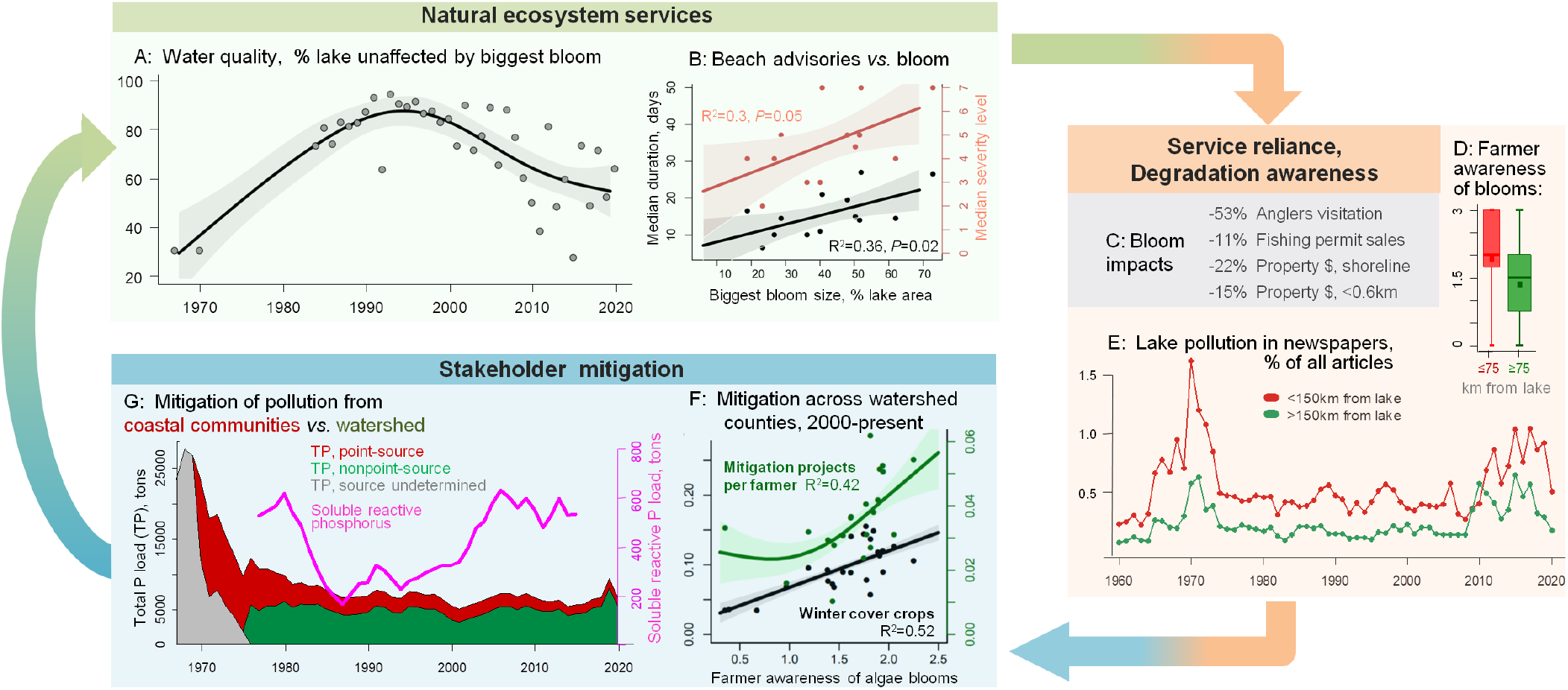
Water quality degradation in western Lake Erie (A,B) affects reliance on lake (C) and temporarily elicits widespread public attention (D,E); reliance and attention correspond to 1970s mitigation of coastal pollution (G) and current efforts to mitigate watershed pollution (F). (A) Percent of western Erie not affected by the largest measured bloom in each year. (B) Years with larger maximum bloom size have longer (black) and more severe (maroon) beach use advisories (medians across 66 Ohio beaches). (C) Persistent blooms reduce recreational lake use and property values. (E) High-impact blooms correspond to brief periods of high newspaper coverage, which is greater in communities closer (red) than farther (green) from the lake, a pattern reflected in stakeholder (farmer) awareness of blooms (D). (F) Farmer awareness correlates with greater farmer participation in government mitigation grants (green) and greater proportion of farmland covered by winter or perennial crops (black) across counties in the Maumee River watershed (points). (G) 1970s mitigation diminished coastal point-source pollution (red), but remaining pollution from watershed (green) underlies contemporary blooms as climatic changes have increased nutrient solubility (pink line; all values are 5-year running averages). In (A,B,F) lines are best-fitting splines and shaded areas represent 95% confidence intervals; see *Methods* for details and data sources.

In the 1960s, western Lake Erie faced intensive, harmful algal blooms due to point-source pollution and runoff from coastal urban centers (28). The resultant shortages in clean drinking water, limited beach use, and declining fisheries heavily impacted coastal communities (18). Reflecting these trends in pollution and reliance, public awareness of water quality issues was (1) higher among communities in close proximity to the lake and (2) peaked in 1969-1975 (Fig. 4e). Partly in response to this attention, the 1972 US Clean Water Act mandated mitigation of the greatest impacts (e.g., municipal sewage facilities, factories, etc). These measures rapidly reduced point-source pollutant loading by stakeholders in Lake Erie (Fig. 4g), benefiting coastal communities that also incurred the greatest mitigation costs (18).

Increases in nonpoint-source pollution from agriculture since 1990, in combination with climatic changes, have led to a resurgence of harmful algal blooms over the past decade (29; Fig. 4a) that impact the provision of (Fig. 4b) and reliance on (Fig. 4c) natural services. Similar to what was observed in the 1960s and 1970s, public attention to water quality is higher in coastal communities with greater reliance on the Lake (Fig. 4d, e), and greater stakeholder awareness corresponds to greater stakeholder mitigation (Fig. 4f). Despite several state and international policy efforts (30), overall mitigation by individuals (at the farm or field-scale) remains low (31), and nutrient loads continue to rise (32; Fig. 4g). These trends are likely due to two key factors. First, stakeholders in the system further from the coast (predominantly farmers) have been slow to recognize the best practices to mitigate nutrient pollution (i.e., *b_t_* > 0; Wilson et al. 2018). Second, mitigation relies on stakeholder actions from across the 300km-wide watershed, such that attention to water quality is a bigger driver of stakeholder mitigation than reliance on the Lake (i.e., high *f*, low *p sensu* Fig. 2c). Therefore, compared to the 1970s, mitigation in the Lake Erie system now relies to a greater extent on public attention and is more likely to face limited opportunity windows.

Going forward, our model approach suggests a rapid consensus on best mitigation approaches (i.e., reducing *b_t_*) and increasing opportunities for stakeholders to re-evaluate their stance on mitigation (i.e., increasing *κ*) can be similarly effective to direct economic subsidies (i.e., increasing *π_N_*/*c_m_*) in boosting mitigation (Fig. 2c). This is because shorter mitigation delays leave less time for human communities to shift reliance and attention away from water quality. Such acclimation trends are already happening in the system with, for example, $104 million upgrades to Toledo’s water treatment plant (33) and media coverage of algae blooms is declining (Fig. 4e). With mitigation more dependent on public attention, our model dynamics also suggest a slower overall path to mitigation in the current state of the system compared to 1970s reliance-driven mitigation (Fig. 2b vs 2c). Finally, our results warn that protracted management efforts to expand stakeholder mitigation may face a time limit as watershed residents gradually adapt to a green Lake Erie. Thus, a turbid-water regime in this system may arise not via ecological feedbacks present in shallow lakes but through the culmination of ecological and social processes.

## Discussion

### Cross-system management implications

As societies acclimate to degraded ecosystems, our first key finding is that delays in mitigation exponentially prolong the restoration process and ecosystem service shortages - even when government action drives a majority of total mitigation effort. Our second key finding is that mitigation efforts can face very limited time windows to get off the ground before they are doomed to failure by the presence of alternative stable socio-ecological states. This unique result emerges from our focus on socio-ecological feedbacks, whereby delays could reduce the incentives for stakeholders to mitigate if their reliance on the service has been replaced by other options. Recognizing this impetus for early action is particularly critical because an array of factors - slow information spread, improving mitigation technology, and alternative investment opportunities - ubiquitously entice stakeholders to delay mitigation.

The three acclimation processes we examine here (Fig. 1a, 2b-d) commonly arise across conservation issues ranging from resource over-exploitation and pollution to species invasions (Table 1), but differ in their impacts on mitigation efforts. Compared to the case where stakeholders mitigate to maintain their livelihoods or other individual needs (reliance-driven mitigation), we expect longer restoration times and shorter opportunity windows when mitigation is driven by public opinion. This happens because in many systems issue-attention cycles concentrate public pressure in favor of mitigation onto a narrow window of time (leading to Jensen’s inequality), although we note that a fraction of conservation issues appear to experience long periods of high attention (e.g., PCB pollution, acid rain). We also anticipate slower restoration and shorter opportunity windows in the presence of even modestly valuable alternative services that provide an incentive against mitigation that grows over time.

For managers, our results place a premium on monitoring to quickly identify degradation (e.g., intensive pollution or overfishing) and the most viable mitigation strategies. A second requisite ingredient is to rapidly communicate the presence of degradation, mitigation strategies, and the importance of early action to all stakeholders in an ecosystem, highlighting the pivotal importance of public outreach in conservation to trigger the decision making process (34, 35). In addition to outreach efforts after degradation becomes apparent, preemptive outreach helps develop trust and communication forums between stakeholders and conservation agencies (7, 36). These efforts can help bridge the gap between classical ‘top-down’ conservation successes (e.g., point-source pollution) and contemporary mitigation efforts facing the challenge of mobilizing voluntary, collective action by many stakeholders.

In addition to the importance of early mitigation, we find that several societal interventions that can be as impactful as environmental interventions in increasing the chance and pace of restoration. In particular, subsidies that increase the value of natural-state services, taxing alternative services, or concentrating government mitigation onto early phases of degradation might compensate for a delay in stakeholder mitigation (*t_b_*; contours in Fig. 2b-d), a fast rate of degradation (*μ*), or slow ecosystem recovery (*r_R_*).

### Additional societal feedbacks impeding conservation

We emphasize that our results likely under-estimate the importance of early action and the potential for limited time windows by omitting several feedback processes that may operate in specific systems. First, declines in public opinion and reliance we model here may be amplified over the long-term by changes in social memory as people forget the past appearance and services of ecosystems. Such a ‘shifting baselines’ phenomenon appears widespread (37, 38) and may contribute, for example, to a lack of long-term action to address algae blooms in many lakes (39). Moreover, some portion of both psychological and economic acclimation may be irreversible at a certain point, thereby increasing the risk from mitigation delays.

Second, the cost of mitigation may not be constant but instead increase over time with economic path-dependence as sunk costs, economies of scale, and vested interests gradually develop around unsustainable degradation activities (40). For example, water-soluble synthetic fertilizers that increase nutrient runoff can gradually displace organic fertilizers on agricultural markets, and in arid climates such as central USA, agriculture-based economies have developed based on unsustainable rates of water withdrawal from aquifers (41). As with altered-state services (Figs. 1c, 2d), orientation of local economies on environmental degradation (1) creates additional mitigation costs that (2) increase over time when few stakeholders mitigate. Therefore, we expect the development of degradation-based economic activities to form an additional, economic source of path-dependence that limits opportunity windows for restoration.

Finally, multiple individually weak feedbacks acting together can form a stronger cumulative feedback and alternative stable states (5, 42). Here, we show that opportunity windows present before natural service depletion reduces reliance (Fig. 2c) greatly contract in systems where natural service depletion concomitantly gives way to an alternate, lucrative service (Fig. 2d). Additional, ubiquitous feedback processes include (i) ecological feedbacks that can diminish the efficacy of small mitigation efforts (43), (ii) social norm feedbacks that can inhibit the expansion of mitigation to many stakeholders (7), and (iii) perception feedbacks whereby a limited mitigation impact of early-adopting stakeholders raises doubt over the efficacy of mitigation practices. Our study exemplifies how conservation efforts depend on timing (44) and the unique feedback processes that become apparent only in a combined human-environment perspective (13, 27).

### Conclusions

We show that socio-economic and ecological processes intertwine in ubiquitous acclimation feedbacks to not only govern the pace of mitigation but also whether or not restoration succeeds across a broad range of systems. Thus, early action is critical regardless of what feedback loops exist and whether or not they are strong enough to create alternative stable states. Recognizing that acclimation places a premium on early action is especially critical given the array of psychological and economic incentives for stakeholders to “wait and see”. Finally, we point out a close relationship between length of opportunity window and resilience of restored socio-ecological systems, where stakeholders perpetually participate in sustaining ecosystem services. Therefore, the management actions to increase the chance and pace of mitigation that we highlight here may also boost the ability of restored socio-ecological systems to withstand future disturbances emerging in the Anthropocene.

## Materials and Methods

For Lake Erie, we report algae bloom impacts on local economies (Fig. 4c) from (9–11). As an indicator of trends in public attention to water quality (Fig. 4e), we track how often water quality issues are mentioned in 28 Ohio-based newspapers that have the largest number of articles archived on www.newspapers.com. Specifically, for each newspaper and year we find the number of articles mentioning both “Erie” and at least one of “alga*”, “nutrients”, “pollution”, or “water quality”, and then divide this number by the total articles available for the newspaper that year. To control for paper circulation, we then average the frequency of water quality mentions across newspapers located in cities (i) < 150km and (ii) > 150km from the center of western Lake Erie. To verify that public attention affects stakeholders, we also track county-level averages in farmer awareness of Lake Erie algae issues (response scale 1 to 5) in a 2014 survey of farmers in the Maumee River watershed (Fig. 4d; 45). We then compare this metric of farmer awareness to county-level mitigation efforts using best-fit splines in Generalized Additive models in R (package ‘mgcv’; Fig. 4f). We use two metrics of mitigation efforts: (1) the number of government EQUIP grants to reduce nutrient runoff per farmer (2011-2019 averages) and the proportion of all farmland covered by winter commodity or perennial crops in remote-sensed OPTIS surveys (46 2004-2018 averages).

For environmental trends in Lake Erie, we plot maximum algae bloom extent (28, 47, 48) (Fig. 4a) and regress these values against the annual median duration and severity level of advisories posted for 68 Lake Erie beaches in Ohio in 2008-2020 from (49) (Fig. 4b). As an indicator of mitigation trends (Fig. 4g), we used data from (50).

## ACKNOWLEDGMENTS

We thank Daniel Gustafson for access and insight to OPTIS data. This research was supported by the Natural Sciences and Engineering Research Council (MA, CTB), the Dynamics of Coupled Natural and Human Systems Program (BCS-1114934), and the National Socio-Environmental Synthesis Center (SESYNC) under grant number DBI-1052875 (DGW).

## Supporting Information

To analyze the presence of alternative stable states across levels of natural ecosystem service values, we numerically solve model scenarios in Fig. 2d-f starting from two initial conditions: (1) near an undegraded state with *N*_0_ = *R*_0_ = *m*_0_ = 1, *A*_0_ = *D*_0_ = 0.02, *U*_0_ = 0.98, and (2) near a degraded state with *N*_0_ = *R*_0_ = *m*_0_ = 0.02, *A*_0_ = 0.8, *U*_0_ = *D*_0_ = 0. Examining the steady-state natural ecosystem service values that solutions from these initial conditions converge onto, we find that a large range of ecosystems service values lead to alternative stable states where the final system state depends on initial conditions (Fig. S1). The range of conditions leading to alternative stable states expands to greater values of service value in scenarios where the incentives for stakeholder mitigation depend primarily on public opinion (Fig. S1b) or erode due to the presence of alternative services in degraded environments (Fig. S1c).

To analyze how the length of opportunity windows corresponds to ecological resilience of undegraded system states, we quantify opportunity window length and the size of the attraction basin of the undegraded state across ±20% parameter variations in our local sensitivity analysis of reliance-driven mitigation (Fig. 2d). We measure attraction basin size as the proportion of simulations from a set of 1000 initial conditions (randomly drawn from a multivariate uniform distribution with all initial conditions between 0.02 and 1), that converge onto an undegraded socio-ecological state. Under this definition of ecological resilience (51), more resilient system states can recover from a larger magnitude of disturbance (52). We find a saturating relation between window length and ecological resilience (Fig. S2a); as a consequence of this, societal processes in our model (green parameter changes Fig. S2b) can affect ecological resilience as much as environmental processes (dark blue parameter changes).

**Figure S1.**
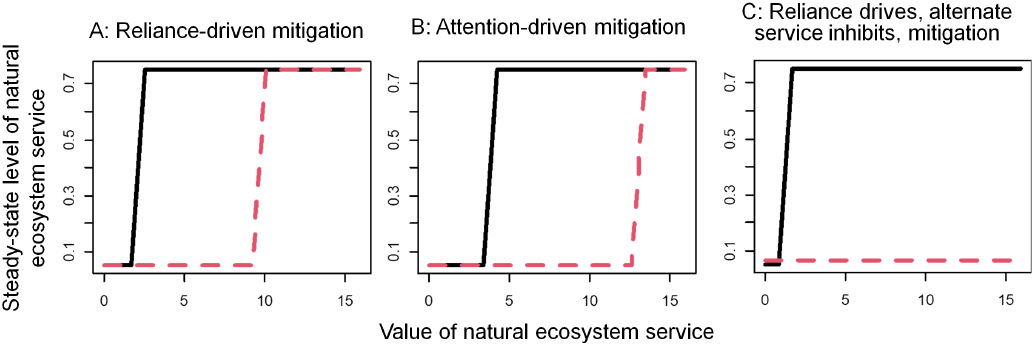
Steady-state levels of natural ecosystem service across levels of natural-state service value *π_N_* in simulations starting from undegraded (black) or degraded (red) initial conditions of the socio-ecological system. Parameter values follow our base scenarios (Fig. 2) with *b_t_* = 0.

**Figure S2.**
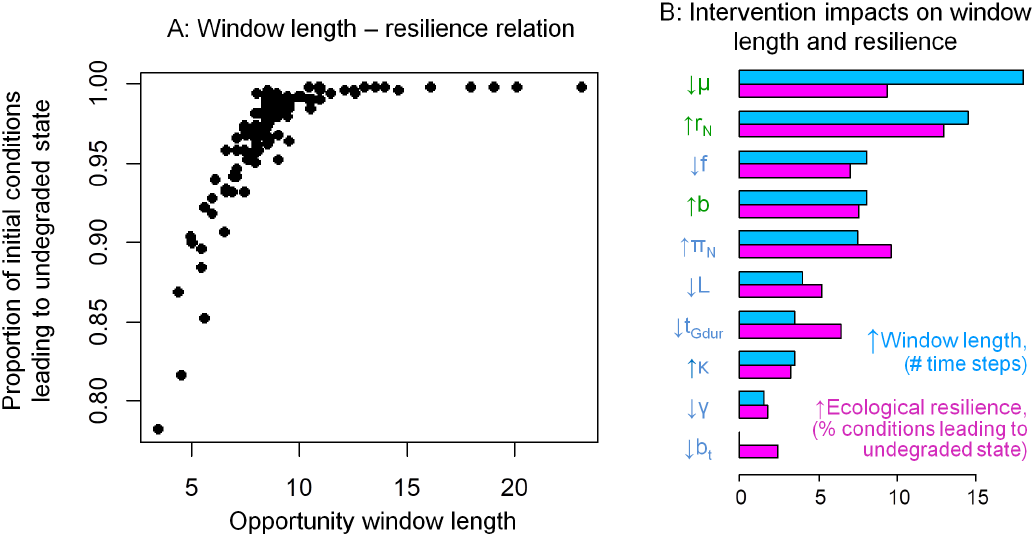
Relation between opportunity window length and ecological resilience (attraction basin size) overall (A) and parsed by changes in each parameter (B). In (B) y-axis labels in dark blue represent changes in societal processes and labels in green represent changes in environmental processes; see Fig. 3c for definitions of each change. Parameter values follow our base reliance-driven mitigation scenario (Fig. 2d).

**Figure S3.**
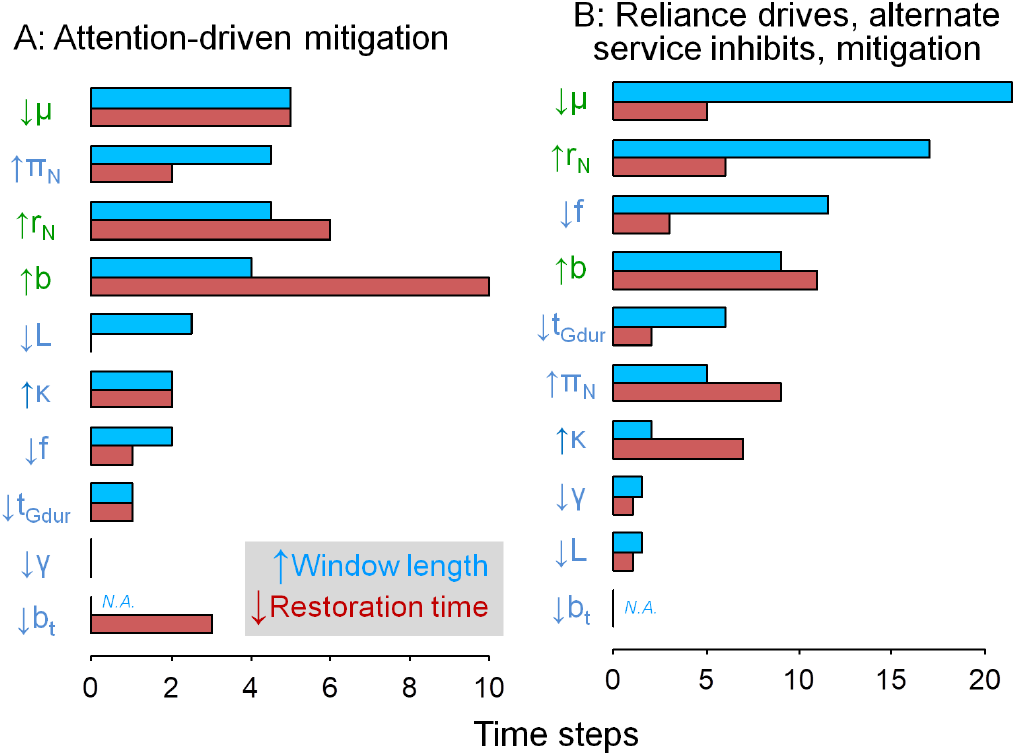
Local sensitivity analyses of the effect of environmental (green) or social (blue) processes in scenarios where mitigation is driven by public attention (A) and where mitigation erodes as alternative services emerge in degraded environments (B). Blue bars represent the decrease in time needed to restore the natural service; red bars represent the increase in opportunity window length (i.e., the maximum time delay *b_t_* that leads to successful restoration). Increases (decreases) represent a change in parameters from 80% to 120% (120% to 80%) of baseline values. Baseline parameter values follow our base scenarios in Fig. 2e and 2f with *b_t_* = 0.

